# Learning and predicting fishing activities from AIS data

**DOI:** 10.1101/2025.04.11.648363

**Authors:** Klaus Johannsen, Xue-Cheng Tai, Gro Fonnes, Junyong You

## Abstract

Marine life significantly impacts our planet by providing essential resources such as food, oxygen, and biodiversity. The surveillance of illegal fishing activities is critical for the sustainable management of marine resources. In this study, we investigate the use of two publicly available data sources to automatically detect fishing activities. One dataset is the Automatic Identification System (AIS) data, the other is the Catch Reports from the Norwegian fishing authorities. The data from fisheries along the Norwegian coast, specifically from vessels with a length of 15 meters or more, covers a period of 1 year and 11 months (January 2022 to November 2023). The AIS data was cleaned by removing duplicates and outliers, then interpolated using piecewise linear interpolation and resampling at a five-minute rate to turn the AIS data into time series of one-hour length. The time series were then tagged using the Catch Report data. The dataset is severely unbalanced with fishing and non-fishing tags. For better learning, the total data was resampled to create 100 balanced bootstrap sets, ensuring equal representation of fishing and non-fishing activities. This process resulted in a benchmark dataset containing about 30 million data points in about 2.5 million time series. We applied a range of classification methods, based on random forests and deep convolution networks. Three types of features were used, related to secant speed, distance to shore, and curvature. We achieved an accuracy ranging from about 92% to 93.7% depending on the features and classification methods used. The predictive capabilities of the classifiers are investigated and significance of the features are studied. The uncertainty of the classification was assessed using the bootstrap sets, providing robust evaluation metrics. Overall, this study demonstrates the effectiveness of leveraging AIS and Catch Report data and advanced data processing techniques for automatic and accurate monitoring of marine fishing activities.

## 1 Introduction

Overfishing poses a significant threat to the sustainability of marine ecosystem fisheries. One of the major contributors to the over-exploitation and decline in marine ecosystem health is Illegal, Unreported, and Unregulated (IUU) fishing. IUU fishing results in an estimated global economic loss ranging from 10 to 23.5 billion annually, with an estimated volume of between 11 and 26 million tons of fish being affected [1]. These figures highlight the critical need for effective management and monitoring strategies.

Norway has placed significant emphasis on sustainable fishing practices, collaborating with international organizations and neighboring countries to curb the negative impacts of overfishing. The country has implemented proactive and comprehensive management strategies, resulting in relatively low levels of IUU fishing activities [2].

The Automatic Identification System (AIS), introduced by the International Maritime Organization (IMO) to enhance maritime safety, has been utilized for collision avoidance, vessel tracking, and monitoring maritime traffic, and is publicly available. AIS broadcasts vessel information that can be received by other ships, ground-based receivers, and satellites [3]. Since May 2014, the AIS has been compulsory for all fishing vessels over 15 meters in length in the EU region [4]. Compared to the Vessel Monitoring System (VMS), AIS offers a higher refresh rate, although it is limited to line-of-sight or VHF ducting propagation. There is a great need to develop new data processing and analysis techniques to extract useful information from AIS data.

Despite these advancements, research on leveraging AIS data for fisheries and its potential for predicting fishing activities from short vessel trajectories remains a challenging field. In this work, we formulate a benchmark problem and propose a method to determine whether a vessel is fishing based on its AIS data from the past hour. The work shows advancements of the current state-of-the-art, pin-points open questions, and opens for further developments.

Several studies have explored various methods and data sources for identifying fishing activities. In [5], the authors identified fishing gear types using VMS data near Indonesia with machine learning techniques. Similarly, in the waters west of Ireland, studies have linked VMS data with Catch Reports to estimate catch per unit effort, demonstrating the practicality of such approaches [6]. Further research has employed AIS data and advanced machine learning algorithms to detect fishing activities, highlighting the potential of AIS data in fisheries monitoring [7–9].

In addition to the above references, several related studies have focused on data analysis techniques for vessel monitoring using AIS data. In [10], they proposed algorithms capable of detecting real-time dark (suspicious or illegal) activities based on vessel movement patterns, aiding in the mitigation of illegal activities such as smuggling and unauthorized fishing. Similarly, [11] developed a real-time maritime anomaly detection framework that specifically targets intentional AIS switch-offs, enabling authorities to identify vessels attempting to evade detection for illicit activities. The work in [12] further analyzed the role of AIS data in detecting dark oil tankers operating in sanctioned maritime environments, particularly in Russian waters.

Data fusion methods have been investigated to enhance vessel detection using AIS data. For example, [13] explored the combination of satellite imagery with AIS signals to improve detection accuracy, particularly in scenarios where vessels switch off their transponders to evade monitoring. In a related study, [14] proposed a data-driven technique to uncover illegal fishing activities by analyzing vessel trajectories and identifying abnormal movement patterns. Similarly, [15] presented a knowledge discovery framework aimed at strengthening maritime intelligence for detecting “dark” vessels, with an emphasis on small, unregulated crafts. Furthermore, [16] surveyed a range of AIS-based anomaly detection approaches, discussing the limitations posed by noisy and incomplete data that can hinder reliable vessel tracking.

In this work, we are mainly interested to learn and predict fishing activities from AIS data. Our tests indicate that predictions based purely on AIS data yield relatively poor results. Therefore, we use Catch Reports to tag AIS data in this study, which shows significant improvements in prediction accuracy in our experiments. We choose to use AIS data for fishing activity detection instead of VMS data for several reasons. The data provides a higher temporal resolution (seconds to minutes) as opposed VMS data (approximately one hour), it is publicly available as opposed to VMS data which is subject to restrictions. Advantages and weak points of our method compared with some afore mentioned literature methods will be point out later in our experiments.

The method proposed in this work, integrating AIS data with Catch Report data, represents an accurate and efficient approach to learn and predict fishing behavior using machine learning techniques. Another contribution of this work lies in identifying relevant features from AIS data. Modern learning techniques, such as deep learning, can automatically extract features from raw data when provided with sufficient ground truth labels. However, this process often requires large, high-quality datasets and substantial computational resources. In scenarios where labeled data are limited as is for our case here, manual feature selection becomes crucial. Through extensive testing, we have identified that numerically determined speed (secant speed), distance to shore, trajectory curvature, and AIS speed are the most important features for identifying fishing activities from AIS data. We show through experiments how these features are contributing to increased prediction accuracy. By using these selected features, we utilize machine learning models to predict fishing activities from a 1-hour trajectory of a vessel based on its AIS reported data.

The organization of this paper is as follows: Section 2 provides a description of the data, including AIS and catch data, and details the preprocessing steps leading to the benchmark problem. Section 3 discusses the classification methods and feature-sets used. Section 4 presents the classification results obtained and discuses the power of the classification methods. Section 5 compares the classifiers and investigates in detail predictive power of the different features. Finally, Section 6 discusses the findings, and Section 7 conclusions are drawn.

## 2 Data, Preprocessing and the Benchmark Dataset

### 2.1 AIS and Catch Report Data

In this work, we will use two public available data sources to predict fishing activities of vessels. One is AIS data and the other is the Catch Report data from fishery authorities. The model derived from this data is able to accurately predict fishing activities of vessels based on AIS data.

AIS data was originally devised for enhancing maritime safety and preventing collisions. AIS is a tracking system that transmits a vessel’s position, AIS speed, course, and other essential information at regular intervals via VHF radio frequencies. This data is captured by other vessels, satellite systems, and terrestrial receivers, enabling comprehensive tracking and monitoring of global maritime activity. The detailed granularity of AIS data allows for in-depth analysis of vessel trajectories, providing insights into maritime patterns including the operational status and movement behaviors of fishing vessels.

Catch Report data, provided by fisheries authorities such as those in Norway, meticulously detail fishing activities, including species caught, quantities, fishing zones, times, and methods. This data ensures sustainable practices, regulatory compliance, and sector transparency. Catch Reports offer insights into fishing effort, productivity, and fish population assessments, informing policies and management. They support sustainable fishing, biodiversity conservation, economic analysis, local communities, seafood traceability, certification, and scientific research. Combined with AIS data, Catch Reports add contextual and outcome-based information, enhancing the understanding of fishing activities and bridging the gap between real-time movement data and empirical results.

For this study, we will use data specific to Norwegian waters. The AIS data was obtained from Norwegian Coastal Administration (Kystverket) and includes only fishing vessels with a length of 15 meters or greater, encompassing about 30 million data points. The temporal coverage extends from January 2022 to November 2023. It includes the entire Norwegian coastline from the Bay of Oslo to the North Cape. The catch data was sourced from Norwegian Directorate of Fisheries [17] and includes more than 700 thousands Catch Reports. In this work, the matching between the AIS data and Catch Report data was performed using the unique vessel ID and corresponding timestamps.

### 2.2 Preprocessing and the Benchmark Dataset

#### 2.2.1 Data Preprocessing

We conducted a series of preprocessing steps on the AIS data to transform it into a benchmark dataset containing time series features and tagged fishing activities. The final dataset comprises approximately 2.5 million time series, each labeled as *fishing* or *non-fishing*. The time series are one hour long, and consist of 12 intervals of 5 minutes each. Each time interval is assigned with features.

The preprocessing begins with AIS data provided by the Norwegian Coastal Administration (Kystverket) and catch data from the Norwegian Directorate of Fisheries (Fiskeridirektoratet). The AIS data is initially formatted as CSV files, and the preprocessing involves various steps to clean and transform this data.

The details of the data processing steps are supplied in Appendix A. The resulting benchmark dataset is stored in `NumPy arrays` and includes time series features, activity tags from Catch Reports and AIS data, bootstrap set indices, and additional metadata.

#### 2.2.2 Data Preparation

After conducting numerous experiments (various different combinations of features used with different classification methods such as random forest, k-nearest neighbor, (convolutional) artificial neural networks), we have concluded that the following four features are the most crucial for predicting fishing activities of vessels

- Secant speed: computed by converting longitude and latitude coordinates into 3D positions, and then determining the secant speed by calculating the norm of the vector of the difference between consecutive positions over time. The secant speed is given in knots.
- Distance to shore: calculated as the distance from the ship’s current position to the nearest shoreline. The distance is given in nautical miles (nmi).
- Curvature: calculated using the method detailed in Appendix B. This feature measures the curvature of the trajectory at a specific position.
- AIS speed: as recorded in the AIS data. This represents the instantaneous speed of the vessel as registered by the vessel’s AIS device, which may significantly differ from the secant speed.

#### 2.2.3 Benchmark Data Structure

The benchmark dataset is provided in the form of `NumPy arrays` with specific filenames and dimensions:

∘ `X ts12.npy`: This file contains the time series data for 2,499,739 instances. Each entry corresponds to a 1-hour vessel trajectory, comprising 12 time intervals and 4 features per interval as stated in §2.2.2. The `NumPy array` has the shape `(2499739, 4, 12)`.
∘ `y_ts12.npy`: This file includes activity tags sourced from Catch Reports as explained in step 8 in Appendix A. The `NumPy array` is shaped as `(2499739,)`.
∘ `y_ais_ts12.npy`: This file contains the fishing activity tags registered in the AIS data. This tag is only used in the experiments in §5.5. A 1-hour time series is tagged *fishing* if more than 50% of the time intervals is registered as *fishing* in the AIS data, otherwise *non-fishing*. For more details, see https://www.navcen.uscg.gov/ais-class-a-reports. As the AIS tags are less accurate and produce poor learning results as shown in Section 5.5, we have used the Catch Report fishing activity tags in our learning experiments in Section 4. The `NumPy array` has the shape `(2499739,)`.
∘ `bidx ts12.npy`: As explained in Appendix A, step 11, most of the time series from `X ts12.npy` are grouped into 100 bootstrap datasets (not all as the balancing of the data imposes some restrictions). The AIS data is imbalanced, the majority having *non-fishing* tags. We split the data into bootstrap sets such that each dataset is balanced with respect to *fishing* and *non-fishing* tags. We will train our models on all bootstrap datasets separately. This file contains bootstrap indicator values *bidx*[*b, i*] = 1, 2 or 3, represented as a `NumPy array` with the shape `(100, 2499739)`. As explained in (1), *bidx*[*b, i*] = 1 indicates that the time series with index *i* ∈ [0, 2499739[in bootstrap set *b* ∈ [0, 100[ will be used as training data. A value of 2 or 3 indicates that it will be used for validation or test instead. Within every bootstrap set we have 14, 500 time series in the training set, 4, 800 in the validation set, and 4, 900 in the test set, corresponding roughly to a 60 − 20 − 20% split of the data.
∘ `ll ts12.npy`: This file holds the longitude and latitude positions of the midpoints of each interval for all the time series in `X ts12.npy`. It is used to calculate secant speed and curvature in the features explained in Appendix A. The `NumPy array` is shaped as `(2499739, 2, 12)`.
∘ `uid ts12.npy`: This file stores unique ship ID for each of the time series, with the `NumPy array` having the shape `(2499739,)`. It is used to derive the fishing activity from the Catch Report data.

Each time series is 1-hour long. It has a unique id (*i* ∈ [0, 2, 499, 739[), being the index to access information in the associated `NumPy arrays` (*X*[*i*], *y*[*i*], *y*_*ais*_[*i*], *ll*[*i*], *uid*[*i*]). For each bootstrap set *b*, the values of *bidx*[*b*] refer to the unique time series id, defining the train-, validation-, and test sets:

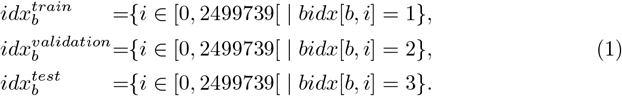

The sets 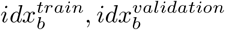, and 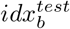 are mutually disjoint with respect to the ships (uids) for each bootstrap set to prevent data leakage. Furthermore, each bootstrap set is balanced with respect to the fishing activity. The classification task for each bootstrap set is to predict the fishing activity (given by the activities derived from the Catch Reports) for each time series. The statistics over the bootstrap sets will be used to assess uncertainty. For comparison with some results reported in the literature [7, 8], we investigate the possibility to model the AIS activity tags as well. It shall be noted that the AIS activity tags are not balanced within the bootstrap sets.

## 3 Methodology

We use the random forest method and the convolutional neural network method [18, 19]. Depending on how many and which types of features are used, we will discuss more specific methods, see below. The training of the methods employs specific hyper-parameter optimization strategies, depending on the methods.

### Random Forest Classifiers

We use three different random forest methods, differing in which and how many features are used:

*RF* (*ū*) : Random forest method using *ū*, the mean of secant speed over the 12 time intervals.

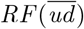: Random forest method using *ū* and 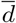 the mean of secant speed and distance to shore over the 12 time intervals.

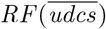: Random forest method using *ū*, 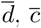,and 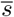,the mean of secant speed, distance to shore, curvature, and AIS speed over the 12 time intervals.

All three random forest methods use time averaged features. To train a particular random forest classifier, hyper-parameter optimization is performed via grid search on the validation set with the following parameter grid:

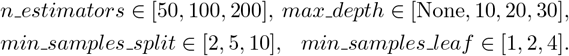

Each of the three methods will be trained on each of the 100 bootstrap data sets (using the train and validation data). We will refer to *RF* (*ū*) trained on the bootstrap set *b* as *rf* (*ū*)_*b*_. The application of *rf* (*ū*)_*b*_ to a feature vector *x* (in this case a 1d-feature vector) is denoted by *rf* (*ū*)_*b*_(*x*). Similar notations will be used for 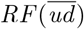 and 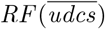,i.e. 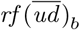 and 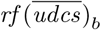, which are applied to 2d- and 4d-feature vectors, respectively. If we referred to the set of classifiers, we omit the subscript, i.e. *rf* (*ū*) etc.

### Convolution Neural Network (CNN) Classifiers

The CNN classification method is using the entire 1-hour time series data, i.e. a feature vector with 48 entries stemming from 4 predictor variables for every of the 12 time intervals of a time series. We used a U-Net type of a Deep Convolution Neural Network method (DCNN) [19]. A good mathematical explanation for Unet was recently given in [20, 21] which has benefited us in selecting the architecture of the Unet used here. We used the following architecture:

- The input for Unet was the dimension 4 features × 12 associated to the 12 time intervals.
- Two or three Convolution layers with ReLU activation and max-pooling were used. This corresponded to the multigrid decomposition of the control variables from the fine and the coarse grids as explained in [20, 21].
- Two or three Convolution layers with ReLU activation and upsampling were used. This corresponded to the multigrid decomposition of the control variables from the the coarse to fine grids as explained in [20, 21].
- A convolution output layer with a scaled softmax activation *S*_*ϵ*_(*x*) = *S*(*x/ϵ*) for classification was used, where 0 indicates *non-fishing* and 1 indicates *fishing*. This corresponded to last layer explained in [20, 21]. *S* is the stadrd softmax function and we take *ϵ* = 0.1 in our tests.
- No skip connection were used.

We denoted the method by *DCNN* (*udcs*). It was trained on each of the 100 bootstrap datasets using the `Adam` optimizer with the `categorial crossentropy` loss function. They were trained on the training sets with a batch size of 600. The training was stopped when the accuracy with the validation dataset reached their best values with respect to training epochs. The *DCNN* (*udcs*) method trained on bootstrap set *b* ∈ [1, 100] is denoted as *dcnn*(*udcs*)_*b*_, the application of it to a 48d feature vector *x* as *dcnn*(*udcs*)_*b*_(*x*).

## 4 Classification Performance

We evaluated the classifiers defined in Section 3 using three key metrics: accuracy, AUC (Area Under the Curve), and confusion matrix.

Let the activity *fishing* be also denoted as positives (P), *non-fishing* as negative (N), and let TP denote the number of true positives, FP the number of false positives, TN the number of true negatives, and FN the number of false negatives. The matrics we will use are detailed in the following.

### Accuracy

This metric measures the proportion of correct predictions among the total number of cases examined. It is a straightforward indicator of overall classifier performance

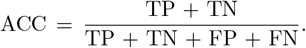

### AUC (Area Under the Curve)

AUC evaluates the performance of the classifier by measuring the area under the ROC (Receiver Operating Characteristic) curve, which plots the true positive rate against the false positive rate at various threshold settings. A higher AUC value indicates a better-performing classifier.

### Confusion Matrix

This matrix provides a detailed breakdown of the classifier’s performance. It allows for a more granular analysis beyond overall accuracy:

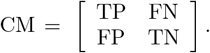

For further details on the metrics, we refer to [22]. We evaluated the classifiers from section 3 using each of the three metrics on each of the bootstrap sets for the train-, validation-, and test sets. In Table 1 we report the results as mean and standard deviation over the bootstrap test sets:

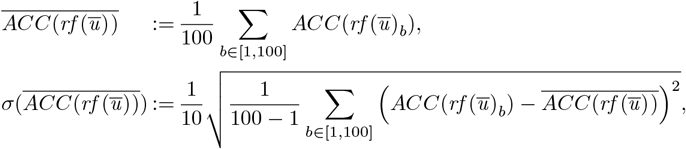

where we used the bias corrected sample standard deviation (cf. 1*/*(100 − 1)) and a reduction by a factor of 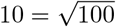 to obtain the standard deviation of the mean. Note that 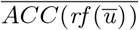 and 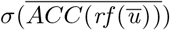 are symbolic notations and don’t have a precise mathematical meaning. In the table this is used as

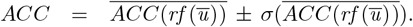

**Table 1.** The performance of the classifiers from Section 3 in terms of accuracy, AUC, and confusion matrix.

| Classifier | ACC [%] | AUC [%] | CM |
| --- | --- | --- | --- |
| $rf(\bar{u})$ | $91.98 \pm 0.11$ | $94.30 \pm 0.11$ | $\begin{bmatrix} 2301 & 149 \\ 244 & 2206 \end{bmatrix} \pm \begin{bmatrix} 5.3 & 5.3 \\ 2.8 & 2.8 \end{bmatrix}$ |
| $rf(\bar{ud})$ | $92.88 \pm 0.11$ | $96.03 \pm 0.10$ | $\begin{bmatrix} 2283 & 167 \\ 182 & 2268 \end{bmatrix} \pm \begin{bmatrix} 5.6 & 5.6 \\ 2.5 & 2.5 \end{bmatrix}$ |
| $rf(\overline{udcs})$ | $93.21 \pm 0.11$ | $96.55 \pm 0.09$ | $\begin{bmatrix} 2291 & 159 \\ 174 & 2276 \end{bmatrix} \pm \begin{bmatrix} 5.5 & 5.5 \\ 2.6 & 2.6 \end{bmatrix}$ |
| $dcnn(udcs)$ | $93.73 \pm 0.12$ | $95.34 \pm 0.11$ | $\begin{bmatrix} 2300 & 150 \\ 157 & 2293 \end{bmatrix} \pm \begin{bmatrix} 5.7 & 5.7 \\ 2.4 & 2.4 \end{bmatrix}$ |

A similar notation is used for AUC and CM, where for CM the construction above is used component-wise. The evaluations are given in Table 1. We can draw the following observations and conclusions from the results:

- The average confusion matrices are balanced with respect to the activity tags (*fishing, non-fishing*), and that holds true also for each bootstrap set separately.
- The accuracies for the classifiers *rf* (*ū*), 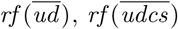, and *dcnn*(*udcs*) are increasing from a mean of 92% for 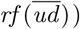 to 93.7% for *dcnn*(*udcs*)).
- Given the values in the table, a two-sample t-test evaluation shows that the differences between the accuracies are statistical significant (p-value is *<* 0.035 in all cases). The same holds true for the AUCs.
- The simplest classifiers, *rf* (*ū*), using the single predictor of the mean secant speed, allows to correctly classify already about a mean of 92% of the cases. This is in sync with the literature [7, 8]. Adding the feature of the mean distance to shore allows for an additional 0.9% of the cases being classified correctly. Adding curvature and AIS speed information allows for an additional improvement by 0.3%. Further improvement by 0.5% to an accuracy of 93.7% is achieved using the complete time series information of all four features with the deep learning classifiers *dcnn*(*udcs*). A further improvement seems not to be possible, see the discussions in the following.
- The three Random Forest-based models 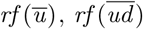, and 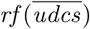exhibit consistently high AUC values, with 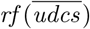 achieving the best AUC of 96.55%. The CNN-based model (*dcnn*(*udcs*)) achieves the highest accuracy (93.73%), yet its AUC (95.34%) is slightly lower than that of the Random Forest models. Upon closer inspection, this drop in AUC appears to be caused by the nature of the DCNN’s output probabilities, which are heavily concentrated near 0 or 1. While this leads to highly confident predictions and strong accuracy, it reduces the model’s ability to rank positive instances above negative ones across the entire range of thresholds, thus affecting the area under the ROC curve. This behavior suggests that although the DCNN is decisive in its classifications, it may not capture nuanced probabilistic distinctions as effectively as the Random Forest models.

## 5 Comparison of the Classifiers

In this section, we compare the classifiers and demonstrate the performance improvements as indicated by statistical metrics. These improvements can be understood through a 3D confusion matrix, the features representing the time series, and their geophysical location. To avoid data leakage, we will only investigate the subset of all time series, which have not been used in the training of the classifiers. It contains about 1.6 million time series and is representative for the entire data set. It is to be preferred over the choice of a particular bootstrap test set, as these contain only 4900 time series. Also, we cannot make a representative choice of one classifier out of the set of all bootstrap classifiers since they do not provide unique predictions, we chose for our investigations the bagging classifier (bootstrap aggregation) derived from the set of bootstrap classifiers. This is a standard choice to create a unique classifier in this setting. We use this setting to compare the classifiers, and assume that this comparison sheds light on the relation of the bootstrap sets of classifiers and eventually on the information content of the features used in these classifiers. We note, that this assumption cannot easily be justified mathematically. We further note that the metrics applied to these classifiers lead to a different accuracy than what was reported in Section 4, due to the imbalance of the dataset with respect to the activity labels (in the data set chosen here for comparison, there are many more *non-fishing* activities than *fishing* activities).

We begin with some definitions and notations. Let *T* denote the set of all 2.5 million time series, *T* ^*^ the set of all timeseries which have not been used in the training of any of the bootstrap classifiers, i.e. not belonging to any of the 100 training or validation sets. *T* ^*^ contains about 1.6 million time series. Let *I* = {*fishing, non-fishing*} be the set of activity labels. We identify *I* with {0, 1}, where *non-fishing* corresponds to 0 and *fishing* corresponds to 1. Further, we define the bagging classifier derived from *rf* (*ū*) by

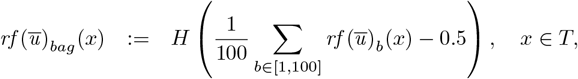

where *H* denotes the Heaviside step function defined by *H*(*x*) = 0, ∀*x* ≤ 0, and *H*(*x*) = 1, ∀*x >* 0, see [23] for details. Similarly we define 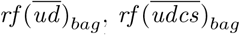, and *dcnn*(*udcs*)_*bag*_. For *x* ∈ *T*, we denote by *y*(*x*) ∈ *I* the activity label (ground truth) of *x*. To compare two classifiers, we use a 3D confusion matrix, which contains the statistical information about both classifiers in relation to each other and the ground truth. In our case of a binary classifier, it is fully specified by 8 numbers. Let *c*_1_ and *c*_2_ be two binary classifiers. Then we define

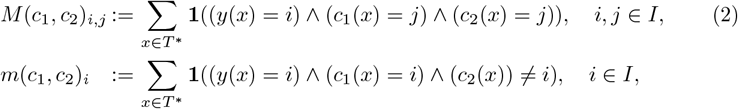

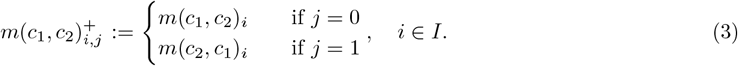

where **1**(*x*) denotes the indicator function, defined by **1**(*False*) = 0, **1**(*True*) = 1 and ∧ the logical AND conjunction. For the classifiers *c*_1_ and *c*_2_, the matrix *M* (*c*_*1*_, *c*_*2*_) corresponds to the standard 2D confusion matrix on the subset of *T* ^*^, where the predictions of the classifiers *c*_1_ and *c*_2_ coincide. The matrix *m*(*c*_1_, *c*_2_)^+^ contains that part of the 3D confusion matrix where the classifiers *c*_1_ and *c*_2_ do not coincide. The first column of *m*(*a, b*)^+^ gives the number of cases where the classification by *c*_1_ is correct and the classification of *c*_2_ is incorrect, for both values of the ground truth. The second column gives the numbers where the roles of *c*_1_ and *c*_2_ are reversed. Both matrices *M* (*c*_1_, *c*_2_) and *m*(*c*_1_, *c*_2_)^+^ together are one possible representation of the 3D confusion matrix, as can be easily verified.

### 5.1 Comparing 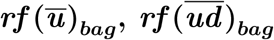

We compare the two classifiers, *rf* (*ū*)_*bag*_ and 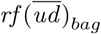, which use one and two time-averaged features, respectively, for the Random Forest method. Evaluating *M* (·, ·), *m*(·, ·)^+^ from (2), (3) results in

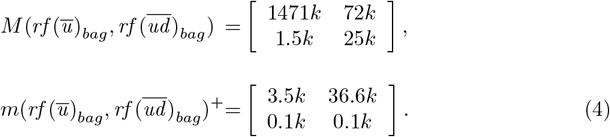

We recall that the first, top rows of *M* (·, ·) and *m*(·,·)^+^ correspond to the ground truth *non-fishing*, the second, bottom row correspond to *fishing*. We first consider 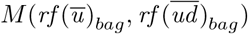.This is the confusion matrix on the subset of *T* ^*^ where both classifiers agree. The dominant entry is the true prediction of ground truth *non-fishing*, showing that the entire dataset is strongly imbalanced towards *non-fishing* activities. Secondly we observe that in 97.5% of the cases the two classifiers agree (∑_*ij*_ *M*_*ij*_*/* |*T*|), i.e. the classifiers are quite equal and the additional information provided by the second feature 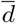 (distance to shore) does change the result only to a small extent. Thirdly, and most interestingly, the major difference between the classifiers occurs for ground truth *non-fishing* in the case where the 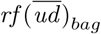-predictions are correct and the *rf* (*ū*)_*bag*_ -predictions are incorrect (*m*_01_*/* ∑ _*ij*_ *m*_*ij*_ ≈ 91%). We concentrate on these 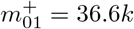 time series. In the following we will call them 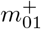-timeseries. In order to understand these time series better we look at their representation in 2D feature space. This is done in Figure 1. In the left image are shown the 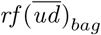-predictions of all time series in *T* ^*^, each represented by a dot. Blue dots represent time series tagged as *non-fishing*, green dots time series tagged as *fishing*. Dominant is the dependence on the secant speed, where most time series with average velocities above 6.3 knots are classified as *non-fishing* and time series with average velocities below 6.3 knots as *fishing*. This corresponds to the predictions of classifier *rf* (*ū*)_*bag*_. The major part, where the two classifiers deviate from each other are ground truth *non-fishing* time series, where the 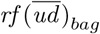-predictions are correct and the *rf* (*ū*)_*bag*_ -predictions are incorrect, the 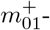 timeseries. These time series are represented with red dots in the right image of Figure 1.

**Fig 1.**
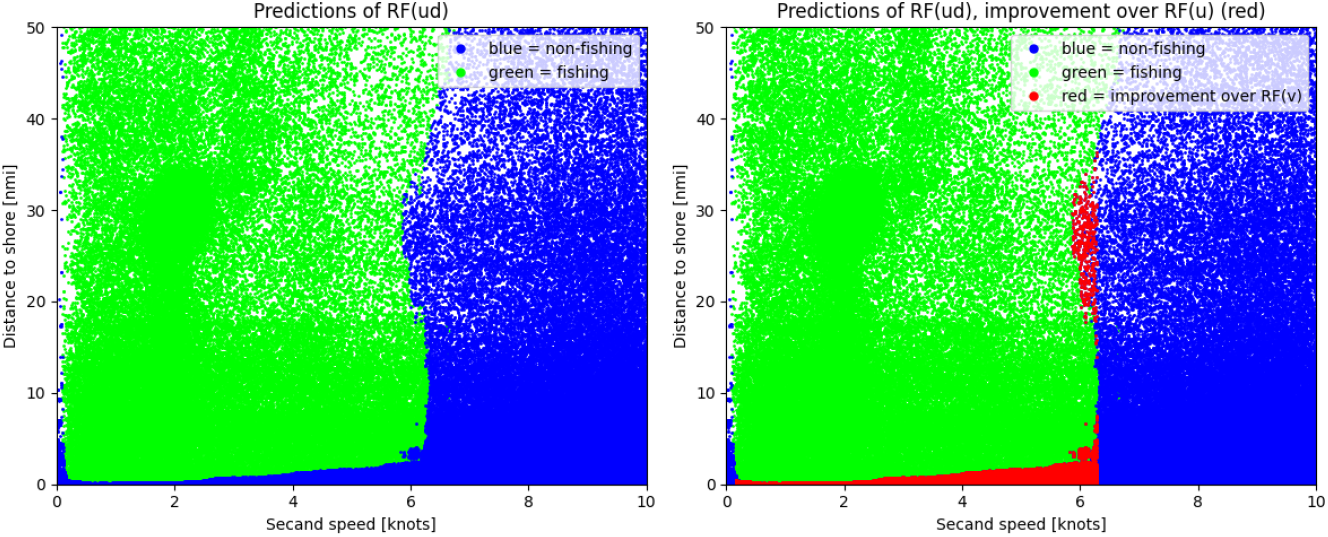
Comparison of *rf* (*ū*)_*bag*_ and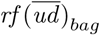. The plot shows the predictions (blue for *non-fishing*, green for *fishing*) of 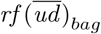 of the 1.6 million time series in *T* ^*^ (left) and the predictions overlaid by the 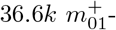 timeseries in red, see eq. (4).

They belong largely to two regions. Most of them lie in the lower band with average secant speed ≤ 6.3 knots and a distance to shore ≤ 10 nmi. These are ship activities close to shore at ‘fishing speed’, which are actually *non-fishing* activities.

The second region is characterized by the upper bound of the ‘fishing speed’, i.e. 6 knots and a distance to shore of 10 to 40 nmi. In geophysical space, the localization of the 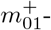 timeseries is shown in Figure 2. The first region corresponds to activities close to shore all along the Norwegian coast. The second region belongs largely to the southern part of the Norwegian Tench, Lofoten, and a region north of the Lofoten. The former corresponds to ship traveling at ‘fishing’-speed. But this lower secant speed is likely kept due to the closeness to shore and not for fishing reasons. The latter region at a distance to shore and secant speeds of around 6 knots is better to be interpreted as *non-fishing*. This would indicate that fishing in that region is rather occurring at slightly lower secant speeds. A statement, which would have to be verified by domain experts, and cannot be done here.

**Fig 2.**
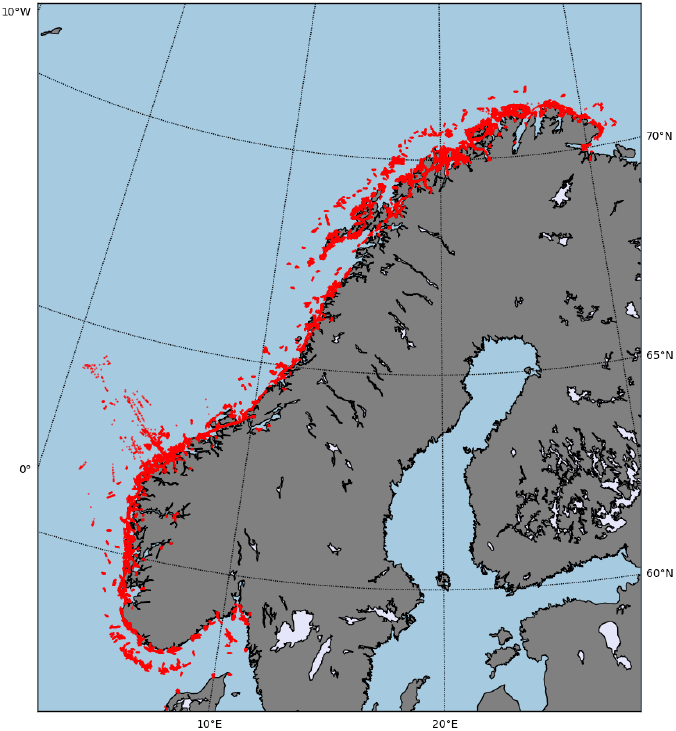
Comparison of *rf* (*ū*)_*bag*_ and 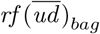. The time series with ground truth *non-fishing*, misclassified by 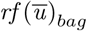 and classified correctly by 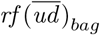 are plotted in red.

### 5.2 Comparing 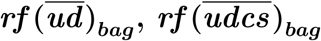

We compare the two classifiers 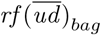and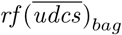, which utilize two and four time-averaged features, respectively, in the Random Forest method. Evaluating *M* (·, ·), *m*(·, ·)^+^ from (2), (3) results in

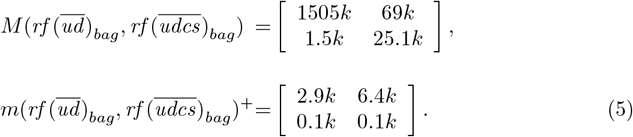

We recall once more that the first, top rows of *M* (·, ·) and *m*(·, ·)^+^ correspond to the ground truth *non-fishing*, the second, bottom row correspond to *fishing*. We first consider 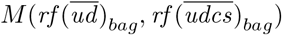.This is the confusion matrix on the subset where both classifiers agree. The dominant entry is again the true prediction of ground truth *non-fishing*. We observe that in 99.4% of the cases the two classifiers agree (∑ _*ij*_ *M*_*ij*_*/*|*T*|), i.e. the classifiers are very equal and the additional information provided by the third and fourth feature 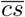 (curvature, AIS speed) does change the result only to a very small extent. The major difference between the classifiers occurs for ground truth *non-fishing* in the case where the 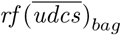-predictions are correct and the 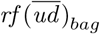 -predictions are incorrect (*m*_01_*/*∑ _i*j*_ *m*_*ij*_ ≈ 68%), the 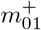-timeseries. We will concentrate on these time series. Note that in this case the effect is less pronounced (the number of cases where 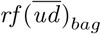-predictions are correct and the 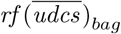-predictions are incorrect is also quite large (2.9*k*)). In order to understand the *m*^+^ -timeseries better we look at their representation in 4D feature space. As points cannot be visualized directly in 4D space, we map the points into 2D subspaces by coordinate selections given as follows. Let 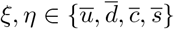 with *ξ* ≠*η*. Then we define 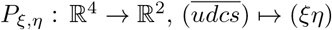,with a slight abuse of notation to simplify formalities.

There are six possible pairings (excluding permutations of the selected coordinates *ξ* and *η*), and we plot the 4D point clouds as six 2D projections. One complication can arise using this representation, which did not occur previously. Time series with different predicted activities can be projected onto (nearly) the same 2D point in the 2D *ξη* plane. A color coding of the predicted classes (*non-fishing, fishing*) as we used before is therefore leading to a non-informative picture. We avoid this by choosing the color of 2D points as a major vote in the following way. We subdivide each 2D plot into 200 × 200 squares and count the occurrences of *non-fishing* and *fishing* tagged time series. The color of the dominant activity tag is chosen for the all points in that square. This procedure blurs the color-class assignments, but the effect is small and an interpretation is well possible. The results for the predictions of 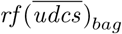 are given in Figure 3. Sets of time series dominantly *non-fishing* are colored in blue, sets of time series dominantly *fishing* are colored in green. On top of that, the 6.4*k* 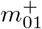-timeseries are plotted in red. They lie close to the decision boundary (the interface region between the blue and green points), indicating that in the ‘core’ regions of *non-fishing* and *fishing* (further away from the decision boundary), the predictions of 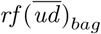 are already good. See also the left image in Figure 1. The 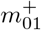 timeseries can be classified better by 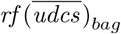 through the usage of curvature information (in addition to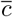, we interpret the relation AIS speed 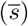 to secant speed (*ū*) as an overall measure of curvature as well). We understand the importance of the curvature information by looking at the top-right and the middel-left images in Figure 3. From the top left, top-right, and middle left images we conclude that the improvements are stemming from two clusters

**Fig 3.**
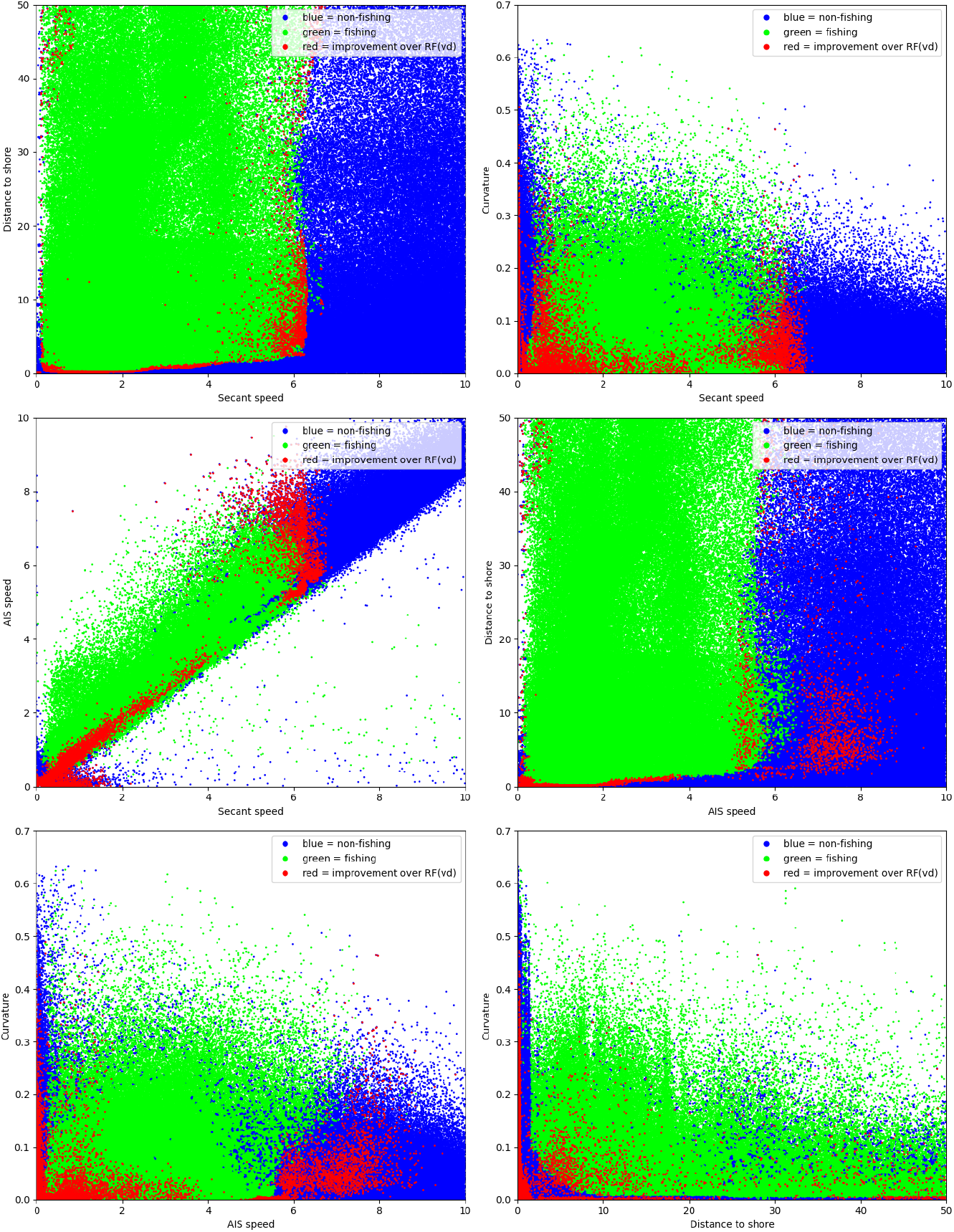
Comparison of 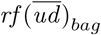 and 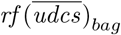.4D points cloud are projected into six 2D points clouds. Blue dots represent time series with a major voting for *non-fishing* activities, green of *fishing* activities, red dots represent time series of *non-fishing* activities, classified correctly by 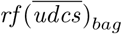 and wrongly by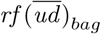.

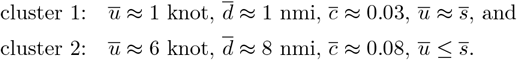

Cluster 1 corresponds to straight, slow cruises close to shore, cluster 2 to curved, faster cruises (max fishing speed), further away from shore. It shows that a combination of secant speed, distance to shore, and curvature was necessary to improve the classification.

In geophysical space, the localization of the 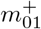 -timeseries are shown in in Figure 5. The clusters cannot be identified in the figure, as curvature information is not directly visible. What can be noticed is a further improvement of *non-fishing* activity classification in the northern regions close to shore (Lofoten and north of Lofoten area) and in the western part of the Norwegian Tench.

### 5.3 Comparing 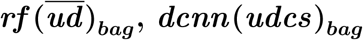

To understand the better performance of *dcnn*(*udcs*)_*bag*_, we relate it to the classifier 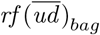 (and not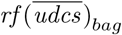). The reason is that, in a sense, *dcnn*(*udcs*)_*bag*_ and 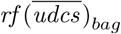 behave similarly, and the results obtained here can be compared to the results from the previous section. We evaluate *M* (·, ·), *m*(·, ·)^+^ from (2), (3):

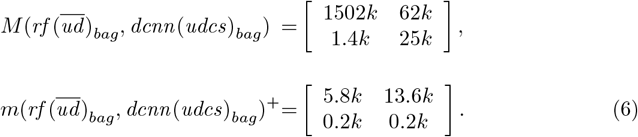

The correct classification of *non-fishing* time series through *dcnn*(*udcs*)_*bag*_, where 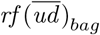 is wrong is significantly larger if compared to the one obtained by *rf* (*udcs*)_*bag*_. These 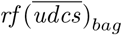 timeseries here are visualized in the same way as in the previous section (Section 5.2). This allows us to directly compare the performance of 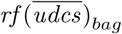 and *dcnn*(*udcs*)_*bag*_. The visualization of the *non-fishing, fishing*, and 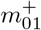 timeseries is given in Figure 4 as before by the blue, green, and red dot, respectively. If we compare Figure 3 and 4, we see that the 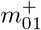 timeseries are in both cases located in the same regions. The difference is that the density of the 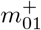-timeseries is higher with the use of *dcnn*(*udcs*)_*bag*_. We conclude that *dcnn*(*udcs*)_*bag*_ is more efficiently able to extract and use information about the curvature. This is also visible when visualizing the 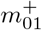 timeseries in geophysical space (Figure 6) and comparing it to improvements obtained by 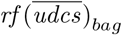(Figure 5). The improvements are largly in the same regions, only the density is higher for *dcnn*(*udcs*)_*bag*_.

**Fig 4.**
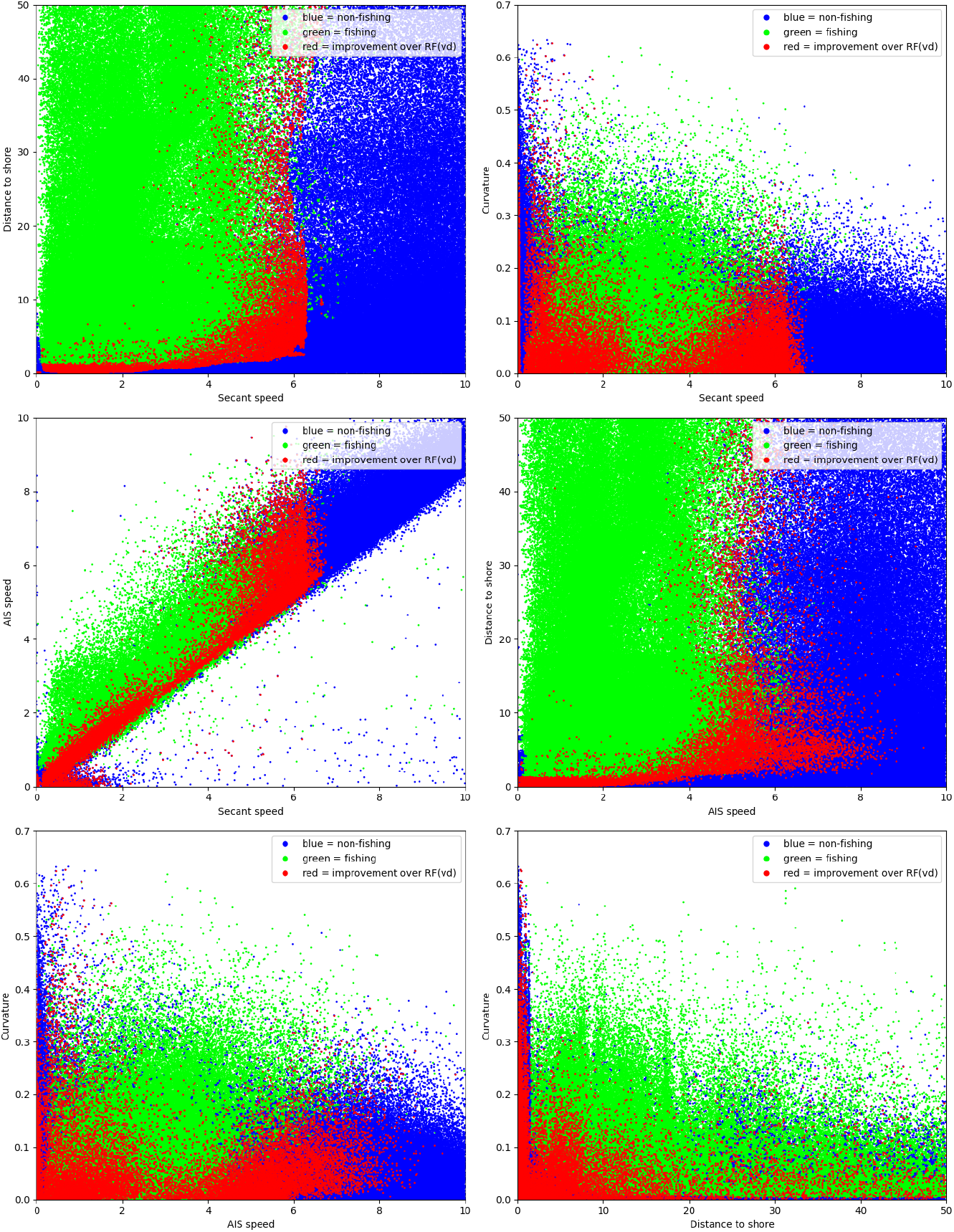
Comparison of 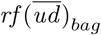 and *dcnn*(*udcs*)_*bag*_. We show the results by six projected 2D points clouds. Blue dots represent time series with a majority of *non-fishing* activities, green of *fishing* activities, red dots time series of *non-fishing* activities, classified correctly by *dcnn*(*udcs*)_*bag*_ and wrongly by 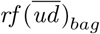 (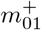timeseries).

**Fig 5.**
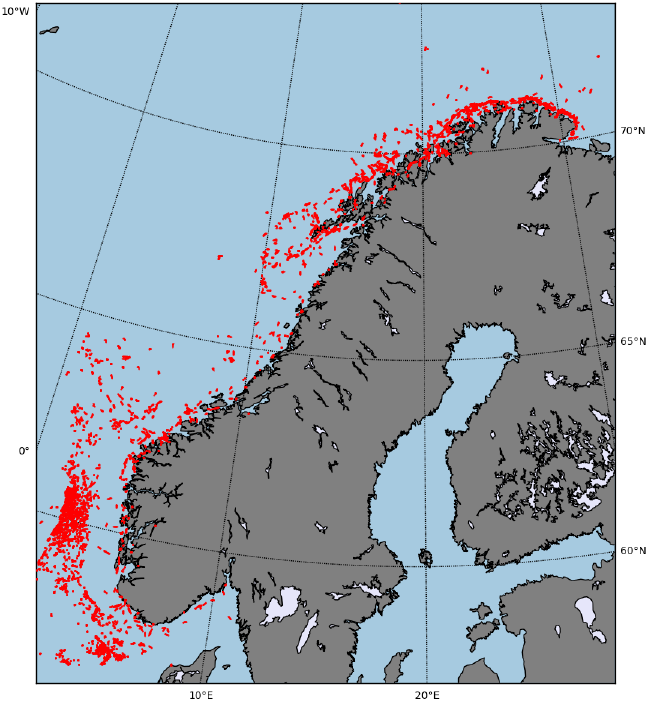
Comparison of 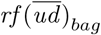 and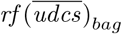. The time series with ground truth *non-fishing*, misclassified by 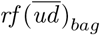 and classified correctly by 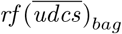 (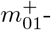 timeseries) are plotted in red.

**Fig 6.**
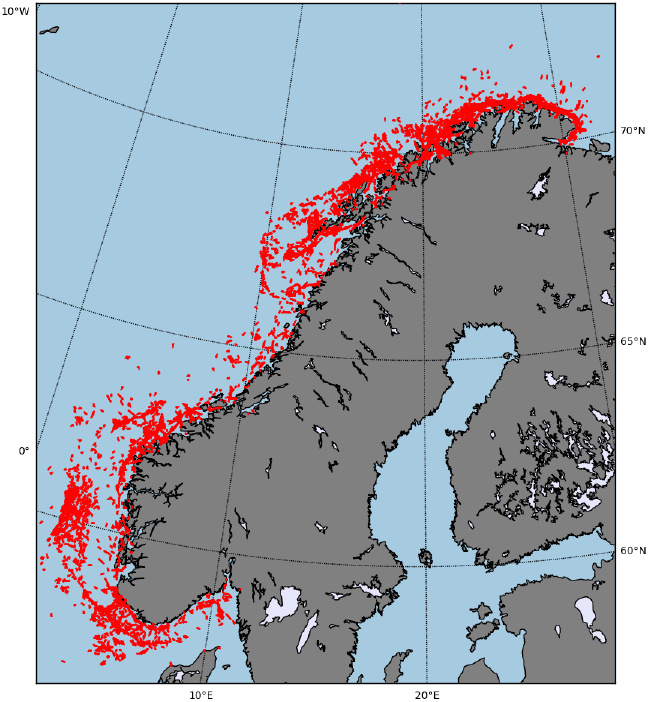
Comparison of 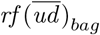 and *dcnn*(*udcs*)_*bag*_. The time series with ground truth *non-fishing*, misclassified by 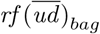 and classified correctly by *dcnn*(*udcs*) _*bag*_ (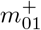timeseries) are plotted in red.

### 5.4 Misclassifications

We obtained a classification accuracy of 91.98% for the *RF* (*ū*) classifier using only the mean secant speed, which can be improved to a 93.71% accuracy using the distance to shore and curvature information, see Table 1. A further improvement seems not to be possible. This raises the question of why this is not possible. The improvements we achieved using curvature indicate that it carries significant information. A reasonable assumption is, if more curvature information was available, e.g. through longer time series, that further improvements would be possible. While an increase of the length of the time series with our software is in principle possible, it alters the time series significantly, so that a direct comparison between shorter and longer time series is not easily possible. On the other hand, the ground truth given by the Catch Report activity tags may actually be wrong, and a prediction might to some extend not be possible. At this stage, we are unable to answer why the accuracy appears to be limited. The explanation could be one or a combination of both reasons given above.

### 5.5 Classification Using AIS Registered Activity Tags

In some cases, the literature reports on the classification of fishing activities by using AIS registered tags, see [7]. Even though the AIS activity tag (navigational status) is supposed to be update by the Officer of the Watch (OOW) manually in a timely manner, there are indications that this is often not done properly, see e.g. [24, 25], which may influence the predictability of the AIS tags. To shed light on the possibility to predict the activities using AIS registered tags given by `y_ais_ts12.npy` (see Section 2), we use the 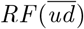 classifier, trained on the AIS tags, and compare the classification accuracy and decision boundary with the results of the same classifier trained to predict the Catch Report tags in `y_ts12.npy`. The results show that AIS tags give very poor results, only slightly better (60.25%) than random predictions, see Table 2. In Figure 7 we show the decision boundary of 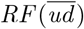 trained on tags `y_ts12.npy` (left) and the decision boundary of 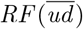 trained on `y_ais_ts12.npy` (right). The decision boundary obtained with Catch Report tags `y_ts12.npy` shows the known pattern of fishing activities associated with vessel velocities below a critical secant speed of about 6 nmi. The decision boundary obtained from the `y_ais_ts12.npy` tags largely classifies the activities as *non-fishing* and assigns *fishing* activities in unrealistic regions of the 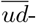 feature space. It shall be noted that a display of the ground truth directly, i.e. the catch report tags and the AIS-tag is not illustrative, as the non-predictable time series of approximately 7% for `y_ts12.npy` and the approximately 40% for `y_ais_ts12.npy` render an interpretation impossible.

**Table 2.**
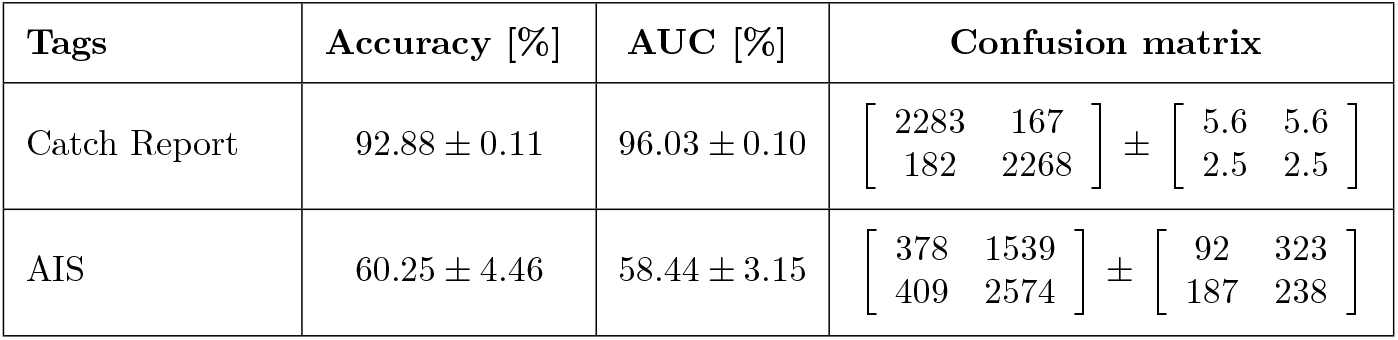
The performance of classifier 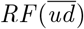 to predict the fishing activities using the Catch Report tags (y ts12.npy) and the AIS registered tags (y ais ts12.npy). The performance is measured in terms of accuracy, AUC, and confusion matrix (the first row / column corresponds to *fishing*, the second to *non-fishing*). The value of the matrices is the mean and the standard derivation of the Confusion matrices.

| Tags | Accuracy [%] | AUC [%] | Confusion matrix |
| --- | --- | --- | --- |
| Catch Report | $92.88 \pm 0.11$ | $96.03 \pm 0.10$ | $\begin{bmatrix} 2283 & 167 \\ 182 & 2268 \end{bmatrix} \pm \begin{bmatrix} 5.6 & 5.6 \\ 2.5 & 2.5 \end{bmatrix}$ |
| AIS | $60.25 \pm 4.46$ | $58.44 \pm 3.15$ | $\begin{bmatrix} 378 & 1539 \\ 409 & 2574 \end{bmatrix} \pm \begin{bmatrix} 92 & 323 \\ 187 & 238 \end{bmatrix}$ |

**Fig 7.**
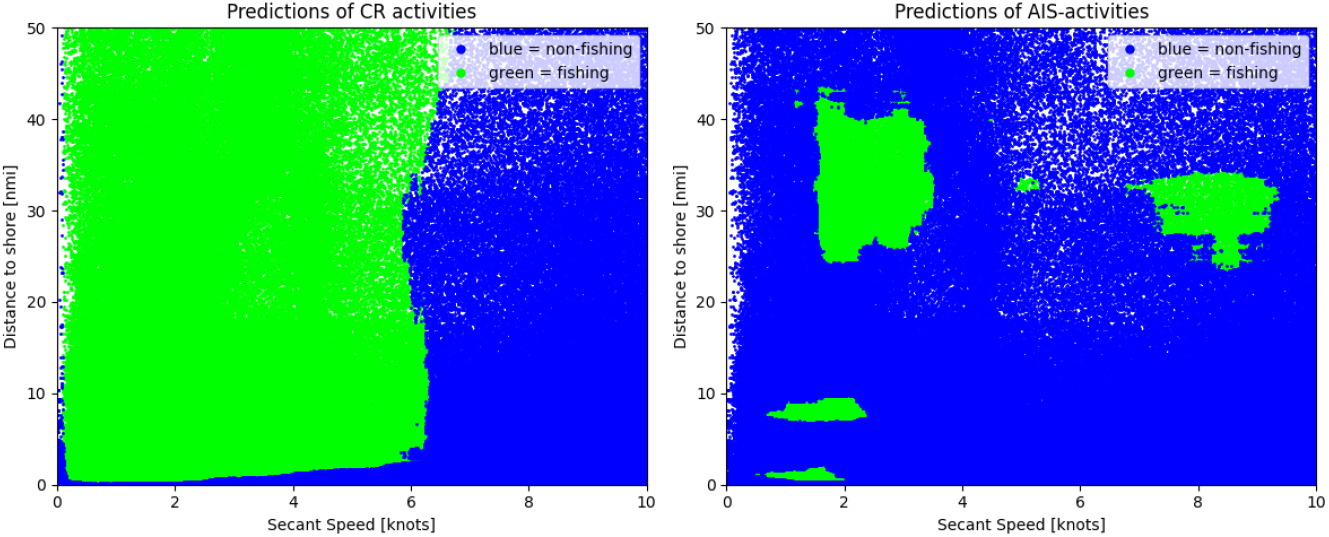
Decision boundary of the 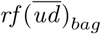 classifier train on Catch Report tags `y_ts12.npy` (left) and AIS registered tags `y_ais_ts12.npy` (right).

## 6 Discussion

In this study, we have addressed the benchmark problem of classifying time series data tagged as fishing or non-fishing activities. Our dataset consisted of 2.5 million time series, each one hour long, providing information on location, secant speed, curvature, and distance to shore. We investigated the performance of different classifiers based on the random forest algorithm and convolution neural networks, with the aim to design the most efficient classifier and to understand the importance of the different features.

Our analysis commenced with the simplest random forest classifier, *RF* (*ū*), which utilized only the mean secant speed as a predictor and achieved an accuracy of 91.98%. This outcome is notable given the simplicity of the feature employed and is consistent with findings from previous studies (Jiang et al., 2016; De Souza et al., 2016). By adding the mean distance to shore as an additional feature (resulting in the 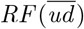 classifier), we observed a slight improvement in accuracy by 0.9%, raising it to 92.88%. This highlighted the significance of a vessel’s proximity to shore as a distinguishing feature for fishing activities.

Including curvature and AIS speed information resulted in an additional accuracy boost of 0.3%, while the most substantial improvement was observed with the deep learning classifier *DCNN* (*udcs*). Utilizing the full time series information of all four features, this classifier achieved an accuracy of 93.71%. The incremental enhancements with the addition of each feature suggest that each one contributes valuable information to the classifier’s predictive capabilities.

Using a 3D confusion matrix enabled a more detailed comparison of classifier performance. For instance, comparing *RF* (*ū*) and 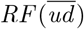 showed that the main improvements happened in the lower range of the feature space, where secant speed was less than 6.5 knots and distance to shore was less than 10 nautical miles. This observation underscored the importance of these thresholds in shaping the decision boundary of the classifier. By comparing the geographical distribution of misclassified time series on the coastal map of Norway, we found that 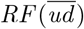 outperformed 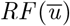 in areas closer to shore, reinforcing the significance of the distance to shore feature.

The *DCNN* (*udcs*) classifier demonstrated superior performance by effectively leveraging the complete set of four features. The time series misclassified by 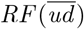 but correctly classified by *DCNN* (*udcs*) were predominantly found in certain areas of the feature space, indicating that the deep learning model excelled at extracting and interpreting complex patterns, particularly from the curvature information.

Despite achieving an accuracy of 93.71%, further improvements appear challenging due to inherent limitations. One such limitation is the amount of curvature information available. Extending the length of the time series could theoretically offer more curvature data, but it would complicate direct comparisons due to significant alterations in the time series. Additionally, inaccuracies in the ground truth data derived from Catch Report activity tags may introduce errors, making certain predictions particularly difficult or impossible. Until more accurate ground truth data or more informative features become available, surpassing the current accuracy limit seems unlikely.

Our investigation into AIS registered fishing activity tags using 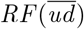yielded an accuracy of only 60.25%, which barely exceeds random predictions. This poor performance highlighted significant limitations in the AIS tags’ ability to provide real-time, accurate information regarding a vessel’s activities. The discrepancy between this and the meaningful classification of Catch Report activities underscores the need for improved tagging and reporting mechanisms for AIS data.

## 7 Conclusion

In conclusion, our study reveals the potential of incorporating multiple features to enhance the accuracy of classifiers for distinguishing fishing activities. Both random forest and deep learning classifiers showed promising results, but the limitations inherent in curvature information and ground truth accuracy present substantial challenges. Future research should aim to extend the time series length without altering its characteristics, improve ground truth accuracy, and explore additional informative features. Moreover, addressing the limitations in AIS data is crucial for enhancing real-time activity predictions, thereby making these classifiers more reliable and applicable in various maritime scenarios.

## Acknowledgments

The authors would like to thank NORCE for making this work possible.

## A Detailed Preprocessing Steps

The preprocessing pipeline includes 11 detailed steps, executed using associated Python scripts (`pp01 xxx` - `pp11 xxx`).

1. Initial Data Processing
  - Read the ship position data, clean it by dropping unnecessary columns and rows.
  - Form trajectories with a maximum time gap of 15 minutes and generate time series with a minimum duration of 6 hours.
  - Carry out linear interpolation and resampling of data at 5-minute intervals.
  - Generate globally unique time series IDs corresponding to non-overlapping 6-hour time series.
2. Ship and Time Series Selection
  - Select 522 ships, each having at least 1,448 time series.
  - Randomly choose 1,448 time series per ship.
  - Save these time series as CSV files, one per ship.
3. Global Time Series Index Creation
  - Create globally unique time series indices starting from 0.
  - Establish mappings between time series IDs and indices.
4. Numpy Array Generation
  - Create numpy arrays with time series data for six variables: longitude, latitude, time, user ID, AIS speed, and nav-status.
  - Each 6-hour-long time series consists of 73 data points sampled at 5-minute intervals.
5. Coordinate Transformation
  - Transform longitude and latitude into 3D spherical coordinates.
6. Distance to shore Calculation
  - Calculate the distance from the ship’s position to the nearest shore.
7. Catch Data Processing.
  - Read the catch data and transform it into the numpy format.
8. Activity Derivation
  - Relate time series and catch data by matching ship id and time.
  - Fishing activity is identified by the corresponding activity in the catch data. Non-fishing tag is assigned in case of absence in the Catch Report data.
9. Feature Calculation
  - Calculate secant speed from successive data points, achieved by transforming positions in longitude/latitude into 3D spherical coordinates and using the time interval length of 5 minutes.
  - Calculate curvature from three successive points using a signed cross-product, Gaussian smoothing, and normalization to ensure that a 90-degree turn over one hour equals 1. For details, see Appendix B.
10. Time Series Segmentation
  - Segment the 6-hour time series into non-overlapping 1-hour time series. The aim was to allow for variable length time series, which couldn’t be exploited in the work.
11. Bootstrap Sampling
  - Create 100 bootstrap samples for training, validation, and testing. The number was chosen to be well over the standard limit 30 from literature 30, see [26].
  - Ensure that the train-validation-test split is approximately 60%-20%-20%, that the data is mutually disjoint in terms of ships, and that it is balanced with respect to fishing activity tags. This results in 14,500 time series for training, 4,800 for validation, and 4,900 for testing in each bootstrap set.
  - Resulting is a utilization of 1,887,752 out of the total 2,499,739 time series across all bootstrap sets.

This detailed description complements the general overview in the main text, providing in-depth information about the data preprocessing steps for those who seek a deeper understanding.

## B Curvature

Given a time series as a sequence of 13 points (endpoints of the intervals) in 3D in units of nautical miles {*x*_*i*_} _*i*=0,…,12_, the curvature {*c*_*i*_} _*i*=1,…,11_ is calculated for every interior point in the following way: Initially, the velocity vectors between consecutive points are computed as Δ*x*_*i*_ = *x*_*i*+1_ − *x*_*i*_. To determine changes in direction, the cross product of consecutive velocity vectors is calculated as *z*_*i*_ = Δ*x*_*i*_ × Δ*x*_*i*+1_. The mean position vector is computed as 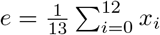, and then normalized to 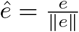. The cross product is projected onto the normalized mean vector to obtain 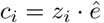. These values are then normalized by the magnitudes of the velocity vectors, *cc*_*i*_ =∥ Δ*x*_*i*_∥ · ∥Δ*x*_*i*+1_ ∥. To ensure smoothness, a Gaussian filter is applied to both the curvature values and their normalizing factors, 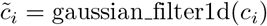and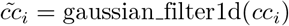. The final curvature for each interior point is then calculated as: 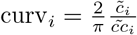. Details of the implementation can be found in the software.

